# Evaluation of the antimicrobial activity and cytotoxicity of different components of natural origin present in essential oils

**DOI:** 10.1101/325639

**Authors:** Sara García-Salinas, Hellen Elizondo, Manuel Arruebo, Gracia Mendoza, Silvia Irusta

**Affiliations:** Department of Chemical Engineering. Aragon Institute of Nanoscience (INA), University of Zaragoza, Campus Río Ebro-Edificio I+D, C/ Poeta Mariano Esquillor S/N, 50018-Zaragoza, Spain.; Networking Research Center on Bioengineering, Biomaterials and Nanomedicine, CIBER-BBN, 28029-Madrid, Spain.; Aragon Health Research Institute (IIS Aragón), 50009 Zaragoza, Spain.

**Author notes:** Corresponding authors: Silvia Irusta, Gracia Mendoza. Both authors contributed equally to the work (Sara García-Salinas, Hellen Elizondo).

**Keywords:** essential oils, bactericida, biofilm, citotoxicity

## Abstract

The antimicrobial action of different components present in essential oils including carvacrol, cinnamaldehyde, thymol, squalene, rosmarinic acid, tyrosol, eugenol and *β*-Caryophyllene against Gram-positive and Gram-negative bacteria is here reported. Planktonic bacteria as well as a model of biofilm forming bacteria were challenged against those components being carvacrol, cinnamaldehyde and thymol the components with the highest antimicrobial action in both different settings. The potential synergy of some of those components against pathogenic bacteria was also analyzed. The antimicrobial mechanism of the different components was analyzed by means of flow cytometry and by electronic and confocal microscopy. Finally, subcytotoxic doses against mammalian cell lines are here reported to highlight the reduced cytotoxicity of those components against eukaryotic cells. Carvacrol, cinnamaldehyde and thymol showed the highest antimicrobial action of all the natural origin compounds tested and lower cytotoxicity against eukaryotic cells than conventional antiseptics such as chlorhexidine. The high inhibition in biofilm forming activity of those components highlight also their demonstrate benefits in reducing pathogenic microorganisms.

**Importance:** The use and misuse of antibiotics has led to the emergence of antibiotic resistance to human and animal pathogens. Compounds from natural sources such as animals, plants, and microorganisms have been proposed as renewed potential antimicrobial alternatives. The comparative antimicrobial action of different components commonly present in essential oils including carvacrol, cinnamaldehyde, thymol, squalene, rosmarinic acid, tyrosol, eugenol and *β*-Caryophyllene against *S. aureus* and *E. coli* is here reported. Carvacrol, cinnamaldehyde and thymol are the components with the highest antimicrobial action. Bacteria membrane disruption represents the bactericidal mechanism attributable to these compounds. In addition, the presence of carvacrol, cinnamaldehyde and thymol hinders *S. aureus* biofilm formation and partially eliminates preformed biofilms. Those components are less toxic to human cells than chlorhexidine.

## 1. Introduction

Since the discovery of penicillin the use of antibiotics, originally developed for human health care, has been extended to animal therapeutics and agriculture (1). The use and misuse of antibiotics has led to the emergence of antibiotic resistance to human and animal pathogens, which is recognized as a serious and global concern because resistance to common bacteria has reached alarming levels in all parts of the world (2). The continued evolution of antimicrobial resistance (AMR) in the hospital is a growing concern because of its potential to endanger the future of antimicrobial drug (3). Even the new generation of antibiotics is becoming virtually ineffective and it is predicted that AMR will cause more deceases than cancer-associated diseases by the middle of the century(4). The discovery of strains resistant to all the antibiotics nowadays available in the clinic have also made an impact on society in general (5). Resistance arises as a consequence of mutations in microbes. Sub-inhibitory antibiotic doses help stepwise selection of resistance and the resulting resistant clones like methicillin-resistant *Staphylococcus aureus, Escherichia coli,* and *Klebsiella* are rapidly disseminated (6). The overall burden of staphylococcal disease caused by methicillin-resistant *S. aureus* (MRSA) strains, is increasing in many countries in both health care and community settings (7). *Escherichia coli*, a common gut bacterium in most mammal species is commonly used to assess the impact of antimicrobials as it carries antibiotic resistance genes. Multiple antimicrobial resistance determinants have been found in *E. coli* on the same plasmid, further facilitating their propagation and co-selection. For instance, the multidrug resistance plasmid IncA/C found in *E. coli*, often encodes for resistance to common antimicrobial agents such as tetracycline, chloramphenicol/florfenicol, streptomycin/spectinomycin, sulfonamides, and extended spectrum β-lactamases, and its spread to pathogenic bacteria may limit antibacterial means to fight infections caused by these bacteria (3).

Because of the emergence of AMR the Center for Disease Control (CDC) has endorsed the need for development of new antibiotics. However, the elaboration of new antibiotics is expensive and time consuming. Meanwhile the genetic plasticity in pathogens results in the development of resistance at a rapid rate (6). Therefore, there is a crucial need for research of new substances with potential to combat resistant strains to minimize their selection. In 2011, academics and industry collaborated on a priority list for approaches to resolve the “antimicrobial-resistance crisis”. Amongst the potential strategies proposed the development of alternatives to antibiotics and the discovery or development of adjuvants were proposed (8). Compounds from natural sources such as animals, plants, and microorganisms have been proposed as renewed potential antimicrobial alternatives (9). Many of them are not new; they have been used to prevent food spoilage since antiquity. Famous seafarers (e.g., Marco Polo) established routes for specie trade and different compounds of natural origin present in species are still being used nowadays to prevent foodborne pathogens showing low levels of antimicrobial (10). Antibiotics by definition have a natural origin (i.e., penicillin is derived from *Penicillium fungi*) and antimicrobials of synthetic origin used to fight against infection are considered as drugs (i.e., isoniazid). However, there are several antibiotics that have multiple mechanisms of action. In this regard, essential oils (EO) are oily aromatic substances extracted from plants with antibacterial, antifungal, insecticidal and antiviral properties. Even when EOs have been distilled for more than 2000 years there is now renewed interest in the antimicrobial properties of phytochemicals and EOs in particular. It is very interesting the demonstrated low levels of induction of antimicrobial resistance towards EOs, that could be related to the fact that these substances do not attack a single specific target but have multiple modes of antibacterial action (1). Antibiotic/drug combinations (i.e., in the treatment of tuberculosis) follow the same principle to reduce the probabilities to develop mutations and consequent drug or antibiotic resistance. Not only antibiotics, but also antiseptics like chlorhexidine, have shown to be able to generate resistance in *Staphylococcus* (11) by mechanisms (mutations in qacA/B gene) which may be common to other microorganisms. However, EO-based compounds are reported as unable to generate antimicrobial resistances in studies involving Gram negative and Gram positive microorganisms subsequently treated with clove, thyme, cinnamon and oregano oils (12–15).

EOs are complex blends of a variety of molecules such as terpenoids, phenol-derived aromatic components and aliphatic components. Their compositions depend on factors such as seasonal variation, climate, plant organ, age, subspecies and even the oil extraction method; consequently, the extracted product can fluctuate in quality, quantity and composition (16). Generally EOs contain about 20-60 components, up to more than 100 single substances, at quite different concentrations; two or three are major components at fairly high concentrations (20-70%) compared to other components that are present only in trace amounts. For example, carvacrol (30%) and thymol (27%) are the major components of the *Origanum* species essential oil. Because of this, in order to have a systematic evaluation of EOs antibacterial activity it is necessary to focus on the study of their main components.

Different components extracted from carvacrol, thymol, eugenol, perillaldehyde and cinnamaldehyde among others have been reported as antibacterial agents (17). However, the reported values for their minimum inhibitory (MICs) and bactericidal (MBCs) concentrations are extremely divergent. For example, the MIC of carvacrol toward *S. aureus* found in the literature ranges from approximately 0.15 mg/mL (18) to 15 mg/mL (19). In some cases the different MICs reported could be attributed to the bacteria strain used. Wang et al. (20) reported a MIC value for carvacrol of 0.31 mg/mL for *S. aureus* ATCC 43300 while using *S. aureus* ATCC 6538 Luz et al. (21) reported a MIC value of 2.5 μL/mL (approximately 2.45 mg/mL) for the same component. But even for the same bacteria strain (ATCC 6538) MIC values for carvacrol of 0.4 mg/mL (22) and 0.015 % v/v (approximately 0.147 mg/mL)(18) can be found in the literature.

Beside the well documented antibacterial action of EO components, there is some evidence corroborating the enhancement in the antimicrobial action of essential oil components used in combination with other antimicrobial agents, both synthetic and natural (23). Thymol and carvacrol were found to have additive antibacterial effect against *S. aureus, E. coli, Salmonella* and *Bacillus cereus*. Ye et al. (24) tested the synergy between cinnamaldehyde and carvacrol in *S. aureus* and *E. coli* among other bacteria and concluded that cinnamaldehyde and carvacrol not only exhibit high antibacterial activities but also have synergistic antimicrobial action against these bacteria. Thymol and carvacrol were found to give an additive effect when tested against *S. aureus* and *P. aeruginosa* (25), besides, thymol combined with carvacrol had a synergistic effect against *S*. Typhimurium (26). One or more synergisms can produce the desired antibacterial effect and reduce the amount of EO components necessary to obtain an efficient bactericidal product.

One interesting application of EO components with antimicrobial activity would be their incorporation into wound dressings since they can prevent or treat wound-associated infections and aid during tissue regeneration (27). Bacterial components have been highlighted as harmful factors during wound healing due to their interference with cell-matrix interactions and due to a reduced inflammatory response they produce. In this regard, *S. aureus* colonize from 30 to 50% of healthy adults and it is able to rapidly infect skin lesions with a consequential inflammatory process (28). *E. coli* is also among the main bacterial species that commonly colonize skin wounds and from this initial colonization, severe problems can occur such as topical infections or even sepsis (29). Within the clinical settings, biofilm formation is a pressing challenge that leads to chronic infections. Prevention of biofilm formation is considered preferable to its removal, since the latter is a very difficult and demanding task, which can cause recontamination problems due to the uncontrolled release of bacterial cells and toxins after their disruption. One of the outstanding antimicrobial properties of many EOs is that they can also be effective even against microbial biofilms (30).

The aim of this work was to shed light on the evaluation of the bactericidal activity and mechanisms of action of EO-present components against *S. aureus* and *E. coli* in planktonic growth. The antibiofilm efficiency and the possibility of synergy between carvacrol (CRV), cinnamaldehyde (CIN) and thymol (THY) were also analyzed. The potential toxicity of those components was also investigated in different cell types, including human dermal fibroblasts, keratinocytes and macrophages.

## 2. Results and discussion

### 2.1. Bactericidal activity against planktonic bacteria

#### 2.1.1. MIC and MBC values

The antibacterial effect of several components present in different essential oils reported as bactericidal, such as carvacrol (31), thymol (32), cinnamaldehyde (33), eugenol (EU) (34), β-caryophyllene (35) and rosmarinic acid (36) were studied. Squalene, a well-known natural antioxidant (37) was also included in the study for comparison. Table 1 and Figure S1 show the MIC and MBC results of the components against *E. coli* and *S. aureus*. The most active compounds were THY, CRV, CIN and eugenol, showing significant differences against the control sample.

**Table 1:**
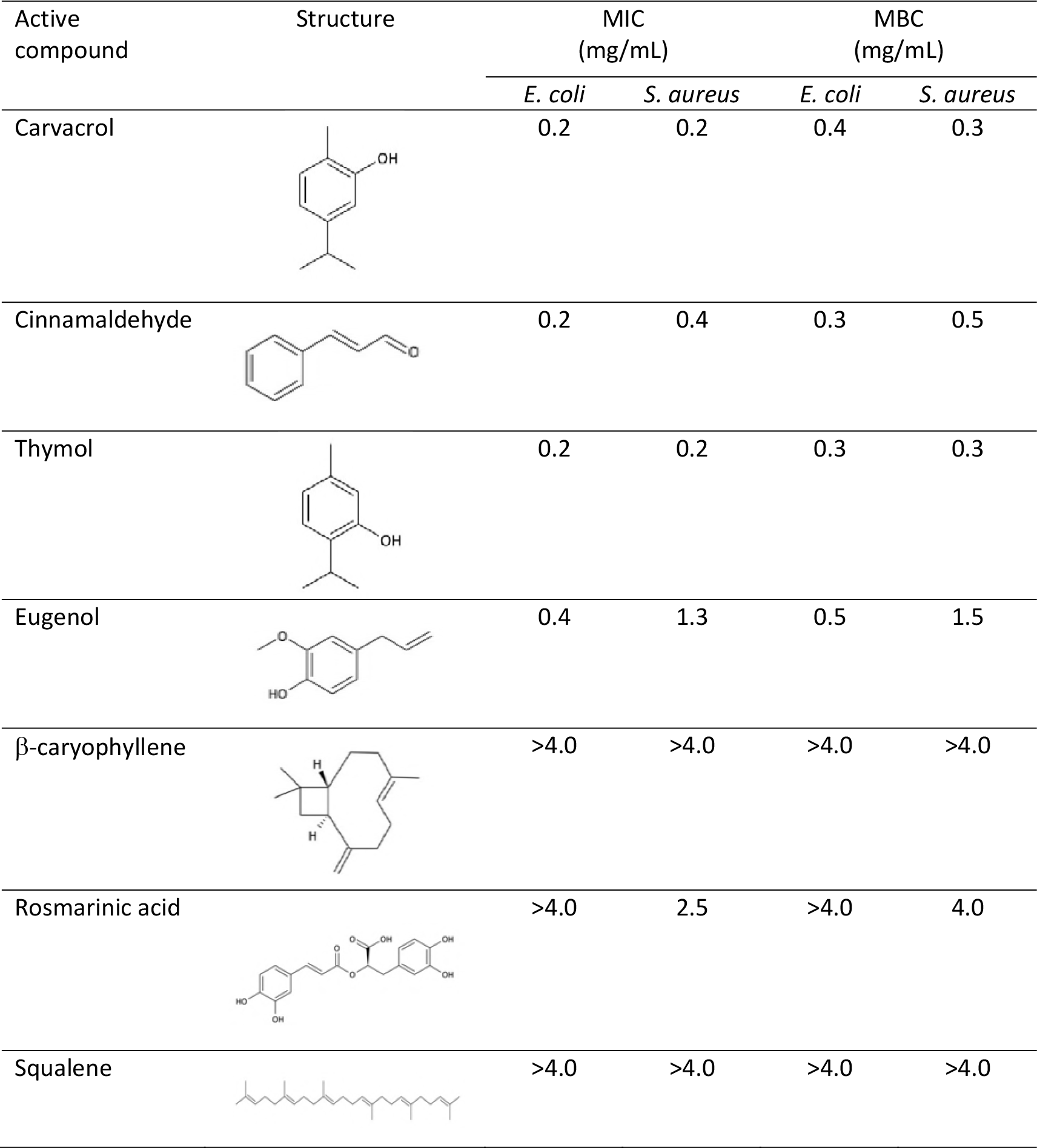
Chemical structure, MIC and MBC values of different essential oil contained compounds. Average of 12 replicas each compound.

| Active compound | Structure | MIC (mg/mL) |  | MBC (mg/mL) |  |
| --- | --- | --- | --- | --- | --- |
|  |  | *E. coli* | *S. aureus* | *E. coli* | *S. aureus* |
| Carvacrol |  | 0.2 | 0.2 | 0.4 | 0.3 |
| Cinnamaldehyde |  | 0.2 | 0.4 | 0.3 | 0.5 |
| Thymol |  | 0.2 | 0.2 | 0.3 | 0.3 |
| Eugenol |  | 0.4 | 1.3 | 0.5 | 1.5 |
| β-caryophyllene | | >4.0 | >4.0 | >4.0 | >4.0 |
| Rosmarinic acid |  | >4.0 | 2.5 | >4.0 | 4.0 |
| Squalene |  | >4.0 | >4.0 | >4.0 | >4.0 |

#### 2.1.2. Bactericidal mechanism

*E. coli* and *S. aureus* exposed during 24 h to THY, CIN and CRV at MIC and MBC concentrations were morphologically examined by SEM (Figure 1). The morphology and microstructure of *S. aureus* before being exposed to any compound can be observed in Figure 1A, having a normal and spherical shape and a well-preserved cell membrane. However, after exposition to carvacrol, cinnamaldehyde and thymol at MIC concentrations for 24 h (Figures 1B-G), the morphology of *S. aureus* cells was distorted. Part of the cell peptidoglycan structure appeared depressed, indicating an initial damage. *S. aureus* exposed to MBC concentrations during 24h became deformed and wrinkled indicating that the intracellular content had leaked out. There was a reduced number of bacteria in the samples and it was hard to find the ones exposed to cinnamaldehyde, probably due to the severe damage to the bacterial peptidoglycan layer and cell membrane and subsequently cell death and detachment from the filter holder. The hydrophobicity of the components present in EOs enables their accumulation in cell membranes disturbing their structures and causing an increase in the permeability allowing intracellular constituents leakage (38). *E. coli* untreated cells were rod shaped, regular, and with intact morphology (Figure 2A) in contrast to MIC-treated cells (Figures 2B-G). SEM images showed morphological alterations and lyses of the outer membrane integrity in cells exposed at MICs. At MBC a complete lysis or seriously damaged cells were observed. The reduction in cell size, length and diameter observed for *S. aureus* in response to the active compound could be reasonably attributed to the leakage of cytosolic fluids outside the cells.

**Figure 1:**
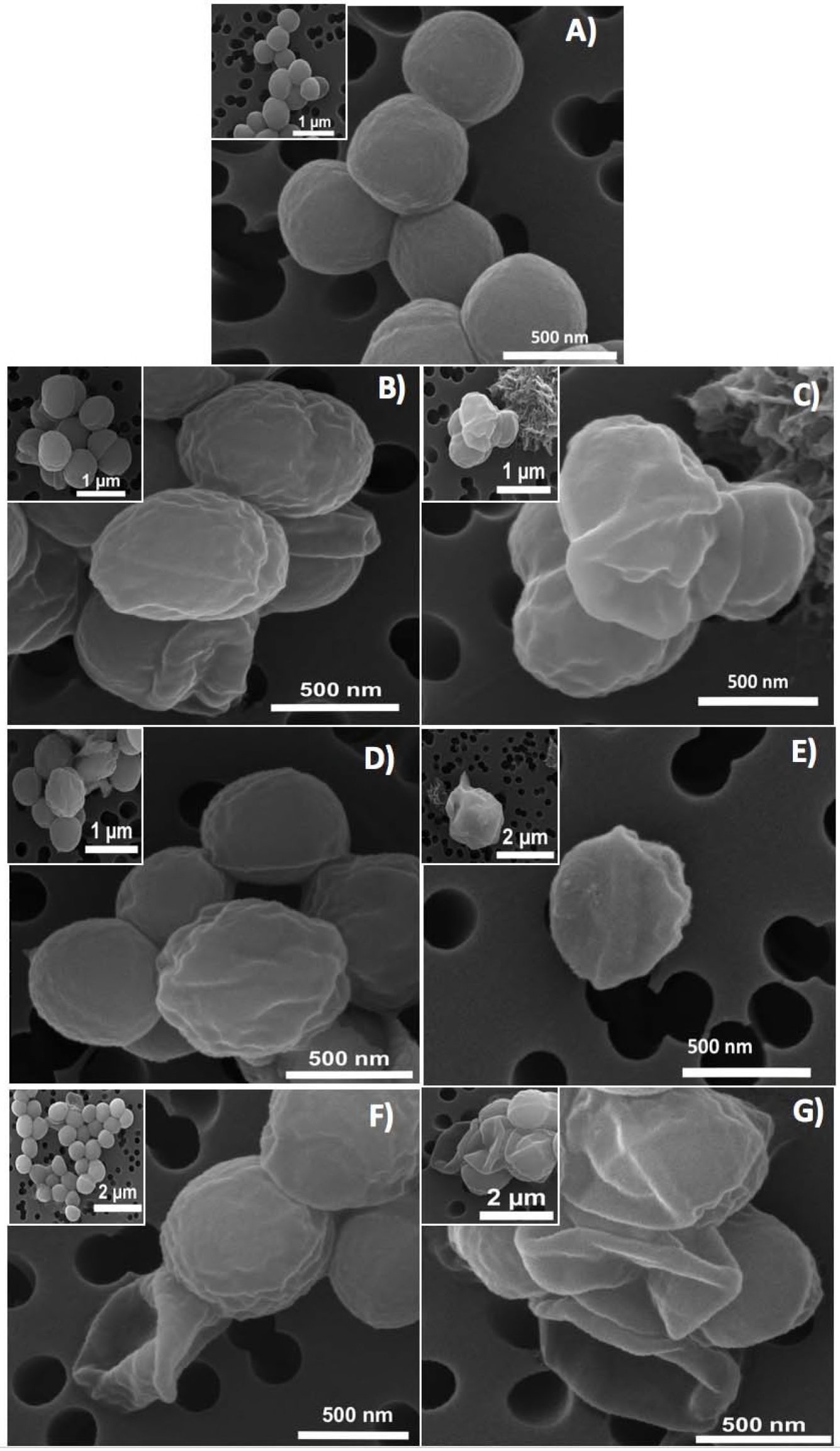
SEM images of *S. aureus A)* untreated (control sample); treated bacteria during 24 h with MIC of B) carvacrol, D) cinnamaldheyde, F) thymol; treated bacteria during 24 h with MBC of C) carvacrol, E) cinnamaldheyde, G) thymol.

**Figure 2:**
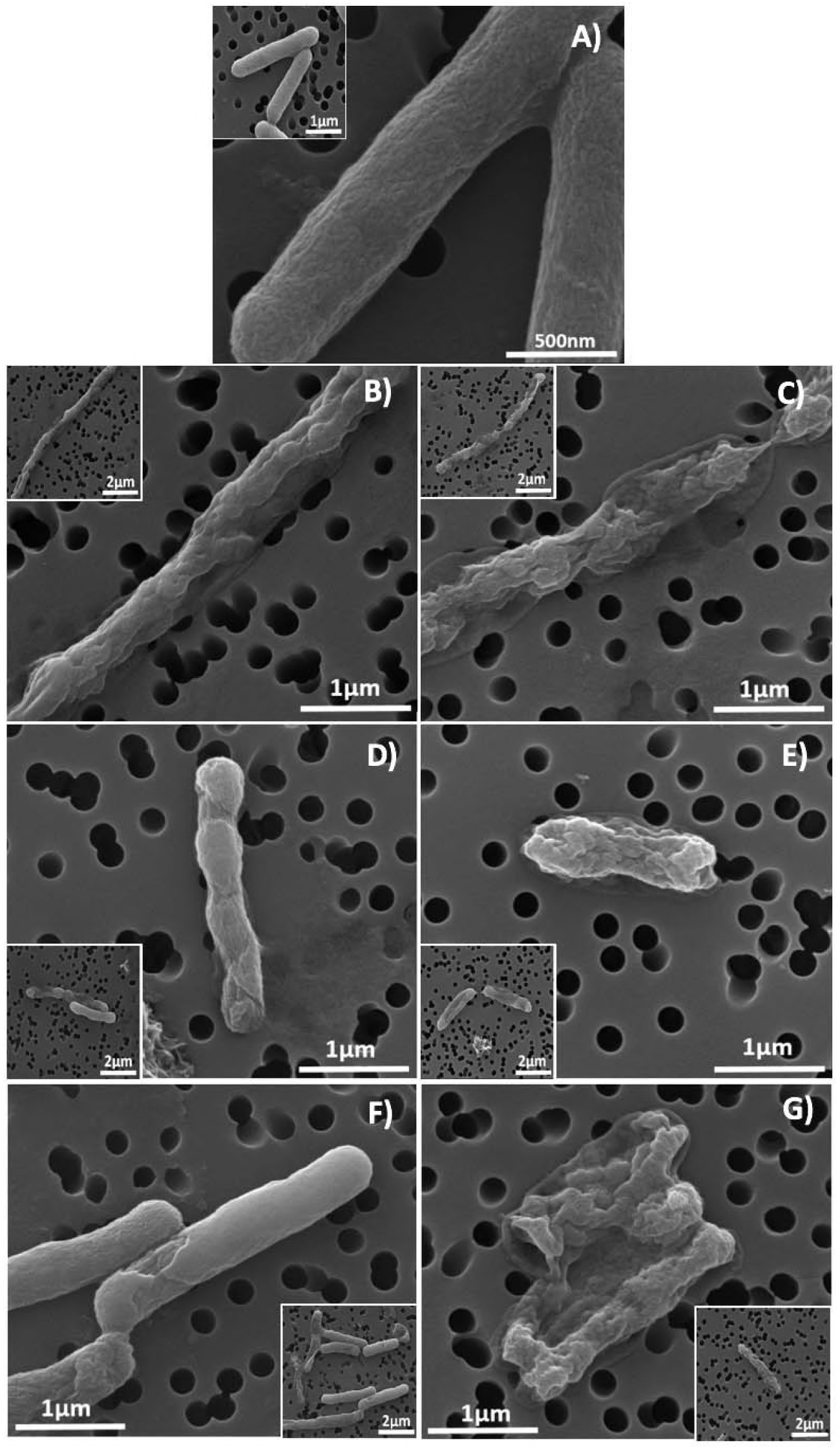
SEM images of *E. coli A)* untreated (control sample); treated bacteria during 24 h with MIC of B) carvacrol, D) cinnamaldheyde, F) thymol; treated bacteria during 24 h with MBC of C) carvacrol, E) cinnamaldheyde, G) thymol.

In order to confirm the bactericidal mechanism of the active compounds present in EOs, flow cytometry and confocal microscopy studies were developed. Flow cytometry histograms (Figure S2) displayed peaks in the range of the negative control (damaged membrane caused by chlorhexidine (39)) when *S. aureus* and *E. coli* were treated with the tested compounds at MBC, which is consistent with cell membrane disruption as previously reported (40). Only cinnamaldehyde treated cells show peaks slightly displaced towards the positive control (undamaged membrane) for both microorganisms suggesting that the involvement of cell membrane disruption in bacteria death was not as clear as SEM images showed. However, confocal microscopy images (Figures 3 and S3) have clearly confirmed membrane damage exerted by the compounds tested when bacteria were incubated with MICs of the compounds, clearly showing red staining related to membrane integrity compromise. Furthermore, in the case of *E. coli* the damaged membrane areas can be clearly distinguished (Figure 3D-F). All these results point out to the bacteria membrane disruption as the bactericidal effect exerted by the compounds present in the EOs.

**Figure 3:**
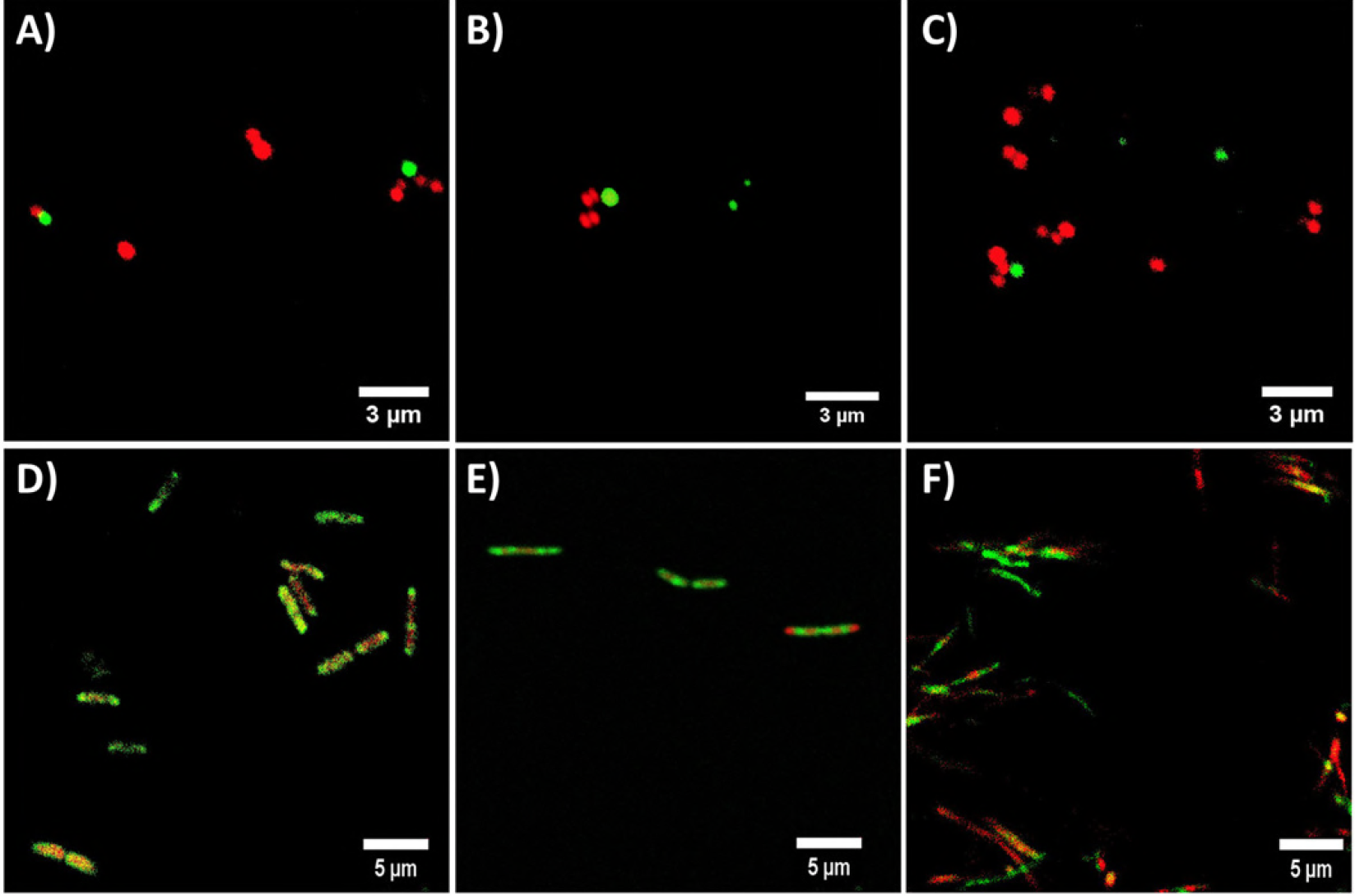
Confocal microscopy images of *S. aureus* (A-C) and *E. coli* (D-F) treated with the MIC of carvacrol (A and D), cinnamaldheyde (B and E), thymol (C and F), stained with the Live/Dead^®^BacLight™ bacterial viability kit. Red staining displays membrane damage.

It is known that phenols, terpenes and aldehydes antibacterial effect is due to their action against the cell cytoplasmic membrane (41). It has been reported that carvacrol and thymol disturb the membrane integrity, increasing the membrane permeability and causing a leakage of protons and potassium finally leading to the loss of membrane potential (42). Di Pascua et al. (41) suggested that the presence of the hydroxyl group in CAR and THY is related to the inactivation of the microbial enzymes. This group would interact with the cell membrane causing leakage of cellular components, a change in fatty acids and phospholipids, and an impairment of the energy metabolism influencing genetic material synthesis. However, some authors have pointed out to different bactericidal mechanisms of action for both compounds due to the different location of the hydroxyl group in their structure affecting cell membrane permeability (43) while others agree with our results showing similar effects for both compounds on bacterial membrane structure (44). It is important to point out that the bactericidal action cannot be related only to the OH group since eugenol having also a hydroxyl group exhibited lower bactericidal effect (Table 1).

According to the literature, the antibacterial mechanism of cinnamaldehyde is not clear. On one hand, it antimicrobial action was attributed to the inhibition of the amino acid decarboxylase activity to bind proteins and no disintegration of the membrane was observed (45). But Nazzaro et al. (43) sustained that like carvacrol, cinnamaldehyde inhibits the generation of adenosine triphosphate from dextrose and disrupts the cell membrane. Our studies would indicate similar trends in MIC and MBC values for CRV, THY and CIN, as well as the disruption of the bacterial surface as target for their activity by three different experimental techniques analyzed.

#### 2.1.3. Synergism

Synergistic interactions between EO active compounds may increase their efficacy as antibacterial agents. For instance, polyethylene films containing a mixture of carvacrol and thymol entrapped within halloysite nanotubes exhibited superior antimicrobial activity against *E. coli* than films containing the individual compounds alone (46). The combination of cinnamaldehyde and carvacrol showed better bactericidal effect compared with the components alone against food-borne bacteria (45). Zhou et al. (26) reported that cinnamaldehyde had a synergistic effect when combined with thymol or carvacrol against *Salmonella* Typhimurium. However, in our case, the FICI values obtained against *S. aureus* (Table 2; Figure S4) indicate that only the CRV-THY combination has an additive effect, while CIN has no interaction with the other compounds tested. The synergistic mechanism between CIN with CRV or with THY was proposed to be caused by the increase in the membrane permeability that enables CIN to be transported into the cell (45). But according to the treated bacteria SEM micrographs (Figures 1 and 2), flow cytometry histograms (Figure S2) and confocal microscopy images (Figures 3 and S3), the effect of the three compounds against *E. coli* is mainly outer membrane disintegration and the morphology of treated *S. aureus* was similar for the three active compounds. THY and CRV were previously found to give an additive antimicrobial effect on *S. aureus* (25) as it was expected since both compounds have almost the same molecular structure (Table 1). It is worth noting that all the FICI values are smaller than 4.0 indicating that there is no antagonism between the tested active compounds.

**Table 2:**
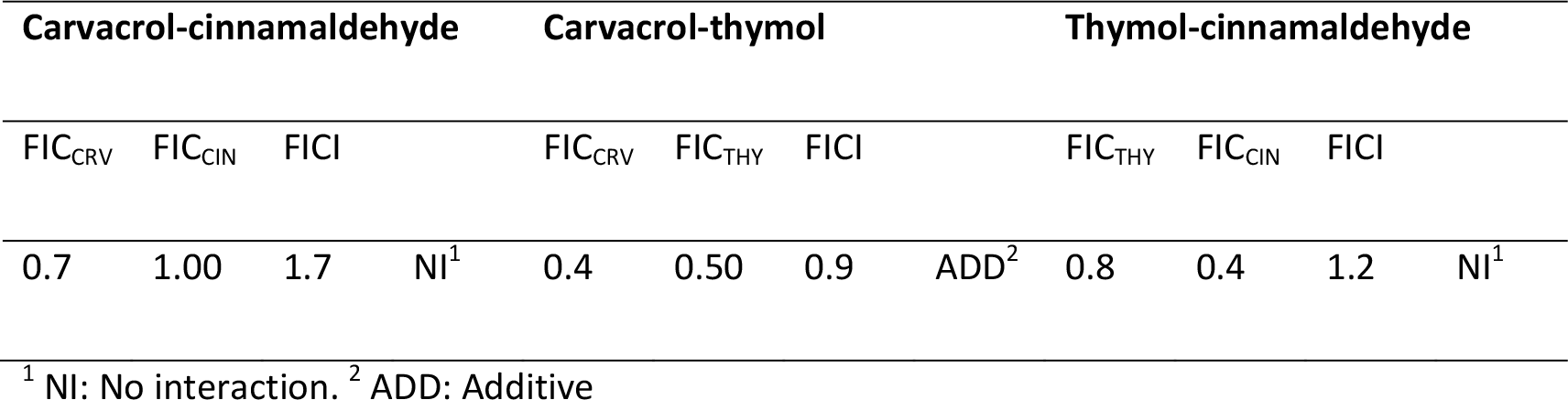
FIC and FICI values for active compounds combination

### 2.2. Antibiofilm activity

*S. aureus* is involved in a wide range of infections that are difficult to treat because, beside the frequent occurrence of antimicrobial-resistant strains, *S. aureus* often reside within biofilms at the infection site (47). Biofilms are communities of microorganisms living at an interphase where they attach to each other through the extracellular polymeric substance also known as the biofilm matrix composed of extracellular DNA, proteins, and polysaccharides. Due to the protection of this matrix, bacteria show up to 1000 times greater tolerance to antibiotics and biocides than their planktonic counterparts (48). Because of this, it is important to find compounds that interfere with the early steps of biofilm formation and slow down its formation rate.

*S. aureus* biofilm formation was observed by calcofluor white staining and by SEM analysis after incubation for 16 h (Figure 4A and B, respectively). The quantification (CFU/mL) carried out after incubation of the formed biofilm (Figure 5A) with the antimicrobial compounds has shown a statistically significant decrease in bacteria growth compared to the untreated biofilms. At 0.5 mg/mL of EO-contained compounds, preformed biofilms showed a reduction in bacteria growth around 2 logs when biofilm was treated with CIN, while CRV and THY exerted a superior decrease (3 logs). The highest tested concentration (1 mg/mL) showed a reduction in CFU/mL higher than 5 logs. As expected, concentrations of the active compounds higher than MIC and even MBC values obtained for planktonic bacteria are needed for biofilm elimination. Our study shows that concentrations higher than 1 mg/mL of any of the compounds tested would be necessary for the total elimination of preformed biofilms.

**Figure 4:**
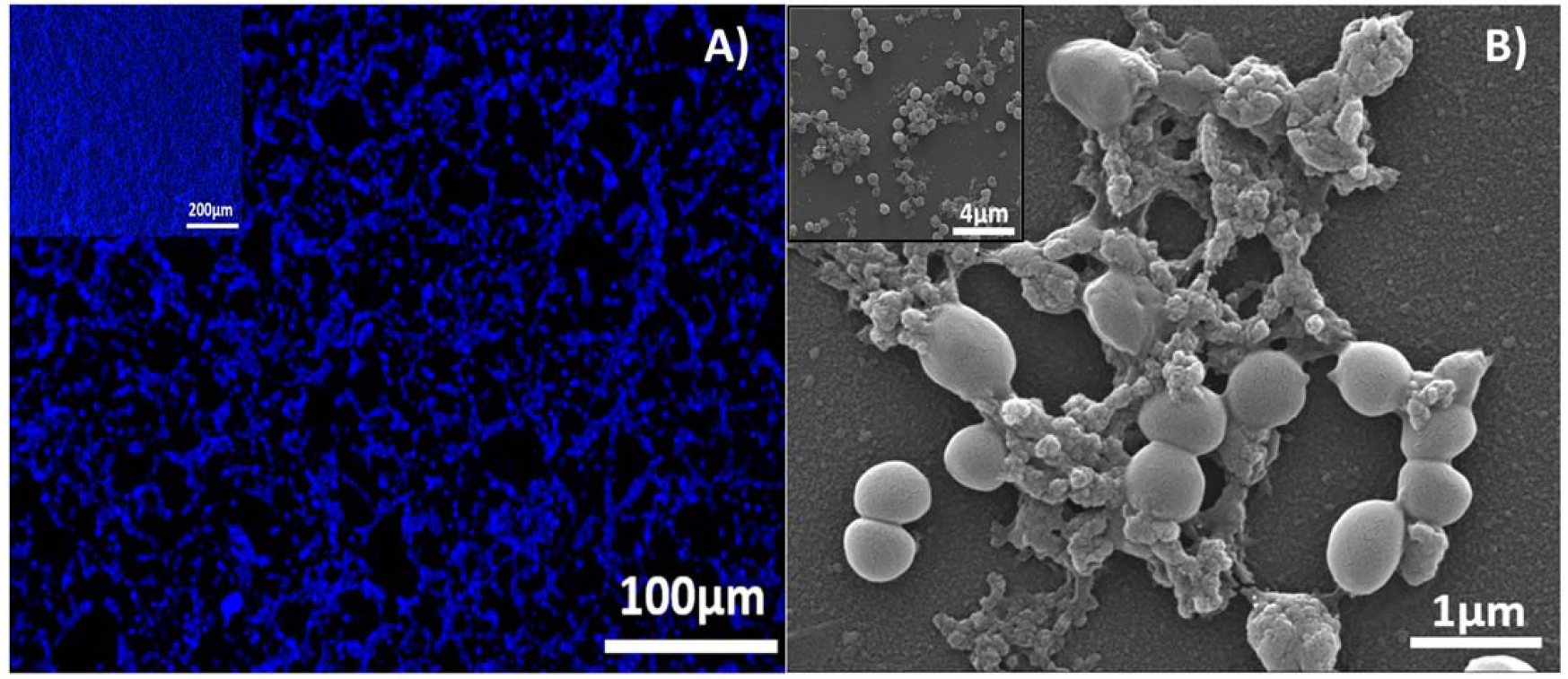
A) Calcofluor staining results and B) SEM image of *S. aureus* biofilm formed after 16h.

**Figure 5:**
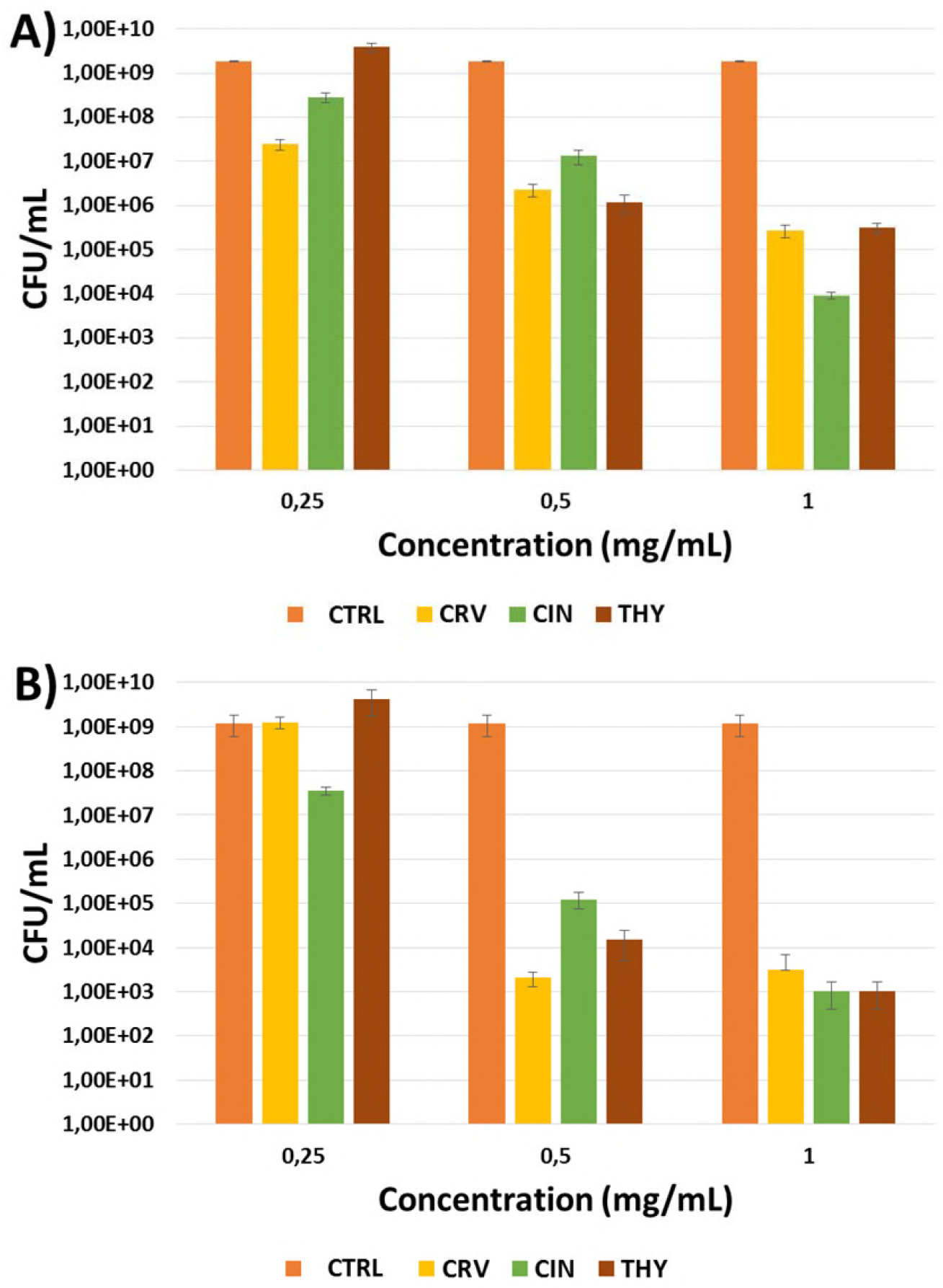
Effect of EOs components at different concentrations on *S. aureus* biofilm: A) elimination of preformed biofilm; B) inhibition of biofilm formation. CTRL=Control sample (not treated biofilm), CRV=biofilm treated with Carvacrol, CIN=biofilm treated with cinnamaldehyde, THY=biofilm treated with thymol.

The addition of EO-contained compounds to the bacteria suspension before biofilm formation hindered this process since there was a significant decrease in the posterior bacterial growth (4 logs for CIN) (Figure 5B). THY and CRV produced even higher reductions of about 5 and 6 logs, respectively. As expected, the concentrations needed to retard biofilm growth and development were higher than the MIC values retrieved for planktonic bacteria (Table 1) but they are in the same range than those reported in the literature (32).

### 2.3. Cytotoxicity

Non-cytotoxicity is required for a wound-healing product since it would be in contact with the infected wound tissue and its neighbouring eukaryotic cells. The cytotoxicity activities of these antimicrobial compounds were investigated using fibroblasts, macrophages and keratinocytes cell lines (Figure 6). Inflammatory cells such as macrophages are generated during wound healing (49), on the other hand, keratinocytes and fibroblasts are part of the epidermis and dermis, respectively. Due to the insolubility of those compounds in aqueous media, those were dispersed thanks to the use of Tween^®^ 80 as described in the materials and methods section.

**Figure 6:**
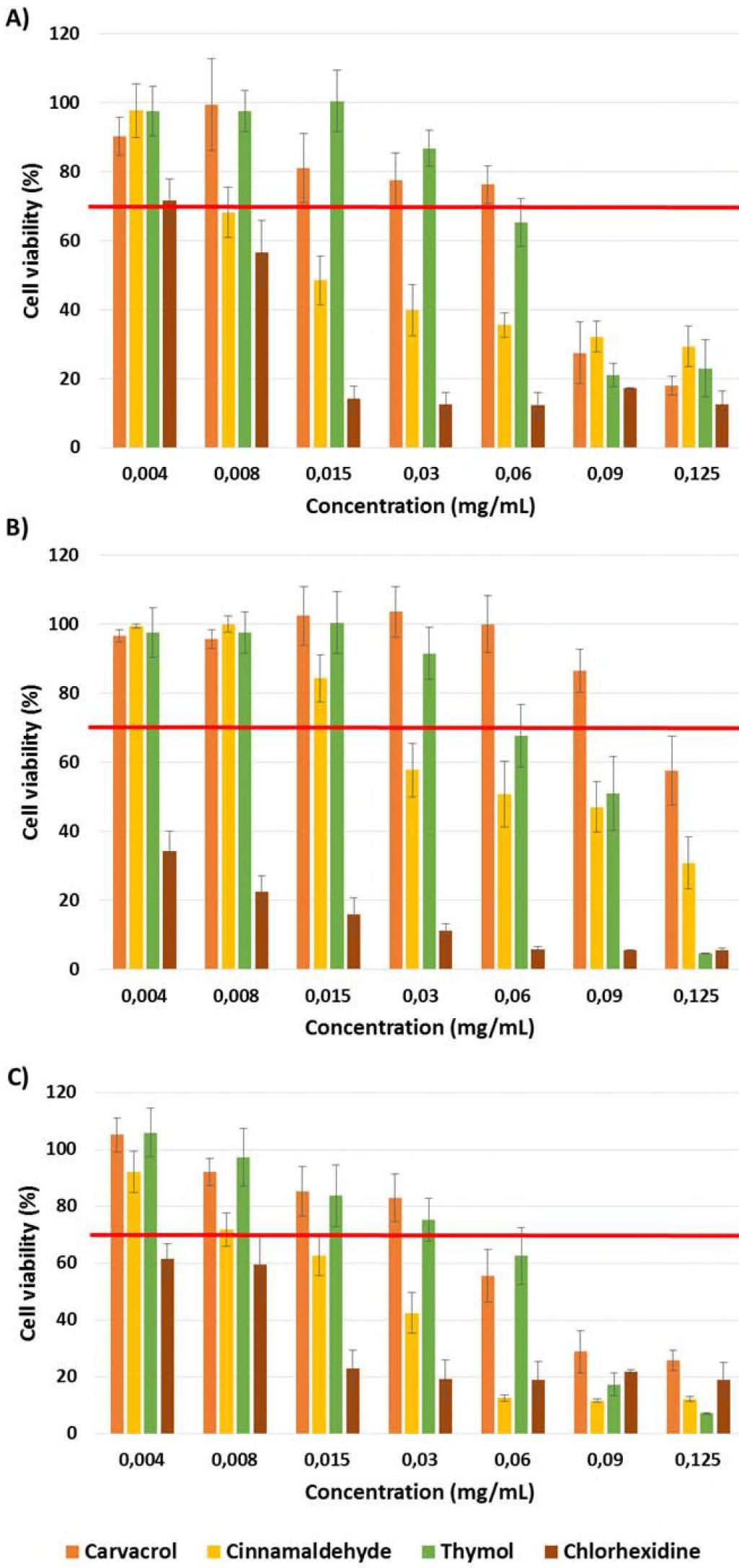
Cell viability after treatment with CRV, CIN, THY and chlorhexidine for 24 h on: A) human dermal fibroblasts; B) macrophages; C) keratinocytes. Control sample (untreated cells) = 100% viability.

Hence, the cytotoxicity of the free compounds would be reduced in absence of Tween^®^ 80 due to their nonpolar character.

CIN was the most cytotoxic chemical of the tested compounds; a dose of 0.030 mg/mL of this compound was enough to reduce the viability of keratinocytes and macrophages below 70% (lowest value established by the ISO 10993-5 to consider a material as non-cytotoxic) and 0. 015 mg/mL affects the fibroblasts viability (Figure 6). Thymol and carvacrol can be considered toxic to fibroblasts at concentrations equal or higher than 0.090 mg/mL and the calculated subcytotoxic doses for keratinocytes were 0.060 and 0.030 mg/mL, respectively. There is also a difference between the effect of THY and CRV on macrophages, since subcytotoxic concentrations were 0.060 and 0.090 mg/mL, respectively. Previous studies have also studied the toxicity of different compounds present in EOs on different human cell lines, such as fibroblasts (50), intestinal cells (51) or different tumor cells (52; 53). Their results show subcytotoxic concentrations for carvacrol and thymol in the same range as ours (^~^500 μM) (51) or higher (50% viability at ^~^5 μg/mL) (50) and also very similar for cinnamaldehyde (^~^10 μg/mL) (53) pointing to apoptosis and membrane damage as key cytotoxic mechanisms.

Chlorhexidine, a typical disinfectant and antiseptic drug used in skin disinfection, was tested for comparison. For the studied concentrations (0.004-0.125mg/mL) chlorhexidine reduces the viability of the three cellular types to 70% or below. Only fibroblasts show viability higher than 70% in presence of chlorohexidine at 0.004 mg/mL. For keratinocytes and macrophages the subcytotoxic concentration was lower than 0.004 mg/mL.

These subcytotoxic values of the tested compounds were lower than the MICs and MBCs retrieved for bacteria, but higher than those obtained with chlorhexidine. In order to reduce bacterial burden in wounds, topical antiseptic agents, among them chlorohexidine gluconate, are usually applied as 2 and 4 v/v % topical solutions (54), concentrations five orders of magnitude higher than the subcytotoxic doses. Therefore, in our study the presence of antibacterial compounds of natural origin in a wound dressing material at MBC concentrations would be only three orders of magnitude higher than the subcytotoxic dose in the worst case scenario demonstrating that those natural origin compounds are less harmful against eukaryotic cells than conventional antiseptics.

Even though the studied compounds showed cytotoxic activity at doses above 0.090 and 0.060 mg/mL, it is important to point out that during the regenerative process in an infected wound the antimicrobial compound at those doses would eradicate both bacteria and eukaryotic cells but while bacteria are removed from the injury eukaryotic cells are continuously arriving to the wound to participate in the regenerative process. Hence, only a small fraction of eukaryotic cells would be eliminated.

## 3. Conclusions

Compounds present in essential oils including carvacrol, cinnamaldehyde and thymol exhibit the highest *in vitro* antimicrobial activities against *E. coli* and *S. aureus* of all the antimicrobials tested. Thymol showed the lowest MBC values (0.3 mg/mL) among all of the compounds tested and was the most effective bactericide against the Gram negative and Gram positive strains evaluated. According to SEM images, flow cytometry and confocal microscopy the bacteria membrane disruption is the bactericidal mechanism attributable to CRV, CIN and THY. There is no antagonism between the tested active compounds, but no synergism was found either; only the CRV-THY combination showed an additive effect. The presence of those compounds at concentrations above 0.5 mg/mL not only hinders *S. aureus* biofilm formation but also partially eliminates preformed biofilms. The subcytotoxic values of the tested EO compounds (in the range 0.015-0.090 mg/mL) are lower than MICs and MBCs for bacteria, but much higher than chlorhexidine doses (0.004 mg/mL). The presence of those antibacterial compounds at MBC concentrations would be only three orders of magnitude higher than the subcytotoxic dose in the worst-case scenario.

## 4. Experimental

### 4.1. Materials

CRV, CIN, THY, Squalene, Rosmarinic acid, Tyrosol, *β*-Caryophyllene, Calcofluor White Stain, phorbol 12-myristate 13-acetate (PMA) and Tween^®^ 80 were purchased from Sigma-Aldrich (St. Louis, MO) while Eugenol was supplied by Acros Organics (Belgium). Tryptone soy broth (TSB) and agar (TSA) were obtained from Conda-Pronadisa (Spain) and *S. aureus* (ATCC 25923) from Ielab (Spain). Regarding cell lines, human dermal fibroblasts were purchased from Lonza (Belgium) and THP1 human monocytes (ATCC TIB-202) from LGC Standards (Spain). High-glucose DMEM (DMEM w/stable glutamine), RPMI 1640 w/stable glutamine and antibiotic-antimycotic (60 μg/mL penicillin, 100 μg/mL streptomycin and 0.25 μg/mL amphotericin B) were supplied by Biowest (France). Cell culture reagents, such as fetal bovine serum (FBS), HEPES, nonessential amino acids, 2-mercaptoethanol 50 mM and sodium pyruvate 100 mM, were obtained from Gibco (UK) and the Blue Cell Viability assay from Abnova (Taiwan).

### 4.2. Bacteria culture

As Gram-negative model *E. coli* S17 strain was used, which was kindly donated by Dr. Jose Antonio Ainsa, while *S. aureus* was evaluated as a Gram-positive model. Both strains were initially grown overnight in TSB at 37 °C under shaking (150 rpm) obtaining, in stationary growth phase, 10^8^-10^9^ colony forming units/mL (CFU/mL). TSA was used for seeding bacteria in an incubator at 37 °C (Memmert, Germany) in order to calculate the bacteria growth (CFU/mL) after treatment with the EO-contained compounds.

### 4.3. Biofilm formation

*S. aureus* was grown overnight in TSB until stationary growth phase was reached. At this point, bacteria were adjusted to 10^7^ CFU/mL and added to a MW96 microplate and incubated at 37 °C for 16 h without shaking. After incubation, culture medium was discarded and biofilms were washed twice with PBS. In order to determine biofilm formation, Calcofluor White Stain was added to each well and incubated 1 min in the dark at room temperature. After incubation, the stain was removed and biofilms were washed twice with PBS. Samples were air-dried in the dark to be further visualized in an inverted fluorescence microscope (Olympus IX81).

To be analyzed by SEM biofilms were grown on sterile glass slides incubated in a *S. aureus* planktonic suspension (10^7^ CFU/mL) at 37 °C for 16 h without shaking. Then, biofilms were washed twice with PBS (0.1 M) and fixed in 2.5% glutaraldehyde for 3 h. Samples were dehydrated through a series of ethanol solutions (30, 50, 70, 80, 90 and 100%; 15 min, twice). Finally, samples were air-dried at room temperature and coated with Pt to allow electronic observation. SEM images were acquired in the energy range of 10-15 keV in a SEM Inspect™ F50 (FEI Co., LMA-INA, Spain).

### 4.4. Antibacterial activity

#### 4.4.1. MIC and MBC determination

Inhibitory and bactericidal concentrations of active EO compounds were tested in two bacteria cultures, *E. coli* and *S. aureus*, following the broth microdilution method. Liquid growth medium containing an inoculum of 10^5^ CFU/mL and serial concentrations of the EO compounds (0.1-4 mg/mL) were used. EO compounds were solubilized in culture medium by adding Tween^®^ 80 (1.5-2% v/v) prior to their serial dilution. Once bacteria suspension was in stationary growth (10^8^-10^9^ CFU/mL) it was further diluted to ^~^10^5^ CFU/mL and added to different concentrations of the antimicrobial agents. Then, samples were incubated for 24 h at 37 °C under shaking (150 rpm). After incubation, bacterial suspensions were diluted in PBS and spot-plated on TSA plates to count colonies after incubation at 37 °C for 24 h. Positive control (untreated bacteria) and negative control (chlorhexidine treated bacteria) samples were also tested.

#### 4.4.2. SEM

Bacteria morphology before and after treatment with EO compounds was analyzed by SEM as we previously reported (55). Briefly, logarithmic growth phase *E. coli* and *S. aureus* bacteria cultures (^~^10^5^ CFU/mL) were treated with the selected EO compounds at MIC and MBC values and incubated overnight at 37 °C. Following incubation, samples were spin-dried at 600g and washed twice in PBS (0.1 M). Bacteria were fixed in 2.5% glutaraldehyde for 90 min and subsequently filtered and dehydrated in ethanol solutions series (30, 50, 70, 80, 90 and 100%; twice for 15 min). Finally, samples were air-dried at room temperature and covered with Pt. SEM micrographs were acquired in a SEM Inspect F50 equipment (FEI Co., LMA-INA, Spain).

#### 4.4.3. Flow cytometry

In order to study the bactericidal mechanism of the different compounds present in EOs *E. coli* and *S. aureus* bacteria samples (10^7^ CFU/mL) were centrifuged at 4400g for 10 min and resuspended in the different compound solutions at MIC and MBC concentrations following the protocols previously described (40; 55). Control groups (not treated and chlorhexidine treated bacteria) were also analyzed. All samples were incubated overnight at 37°C. After propidium iodide (25 μg/mL; Sigma-Aldrich, Germany) addition samples were analyzed by flow cytometry in a Gallios equipment (Beckman Coulter Company, Cell Separation and Cytometry Unit, CIBA, IIS Aragon, Spain).

#### 4.4.4. Confocal microscopy

Live/Dead^®^BacLight™ bacterial viability kit (Molecular Probes, Fisher Scientific, Spain) was used to detect bacteria membrane damage. The methodology is based on the double-staining by SYTO9 and propidium iodide as indicated by the manufacturer. Bacteria samples (10^7^ CFU/mL) treated with the selected EO compounds at MIC were put in contact with the dye mixture for 15 min in the dark at room temperature. Samples were then mounted on slides and visualized by confocal microscopy (Leica TCS SP2 Laser Scanning Confocal Microscope, Microscopy Unit, CIBA, IIS Aragon, Spain). Control samples were also tested as described above.

#### 4.4.5. Synergy studies

The Broth Dilution Checkerboard test was used to evaluate the interaction among the three most promising bactericidal EO compounds determined by the MIC and MBC studies against *S. aureus* as previously reported (56). In brief, a solution containing 4 times the MBC of each compound was prepared. By using a MW96 plate and fresh medium, compound A was diluted two-fold in vertical orientation and compound B was diluted two-fold in horizontal direction. Then, a bacterial suspension (10^6^ CFU/mL, 100 μL) was added and the plate was incubated overnight at 37 °C. After incubation, bacteria growth was determined by the resazurin assay. The Fractional Inhibitory Concentration Index (FICI) of the combination of compounds A and B was calculated according to the following equation:

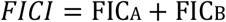

where,

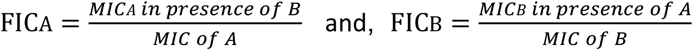

FICI results were classified as synergy (FICI < 0.5), addition (0.5 ≤ FICI ≤ 1), indifference (1 < FIC I ≤ 4) or antagonism (FICI > 4), as previously described (57).

#### 4.4.6. Biofilm disruption

The effects of EO compounds (0.25-1 mg/mL) to prevent the formation of biofilm and to disrupt an already formed *S. aureus* biofilm were studied. EO compounds were added to preformed biofilms and samples were incubated for 24 h at 37 °C without shaking. After incubation, biofilms were disrupted by sonication (15 min, 200 W; Ultrasons, JP Selecta, Spain). Samples were then diluted and seeded onto agar plates to count the viable colonies grown after 24 h of incubation at 37 °C.

To study the effects of the compounds present in the EOs on biofilm formation, those were added to bacterial suspensions (10^7^ CFU/mL) in a MW96 microplate and incubated for 16 h at 37 °C without shaking. After incubation, planktonic cells were removed by washing them twice with PBS. Biofilm samples were then sonicated as described above and serially diluted to be further plated on agar. Viable bacteria (CFU/ mL) were counted after 24 h of incubation at 37 °C.

### 4.5. Cell culture and cytotoxicity assays

Human dermal fibroblasts, human epidermal keratinocytes (HaCaT), kindly donated by Dr. Pilar Martin-Duque, and THP1 human monocytes were used to evaluate the cytotoxic effects of EO-based compounds.

Fibroblasts and HaCaT were routinely grown in high-glucose DMEM supplemented with 10% FBS and antibiotic-antimycotic. Monocytes were cultured in RPMI 1640 supplemented with 10% FBS, 1% HEPES, 1% nonessential amino acids, 0.1% 2-mercaptoethanol 50 mM, 1% sodium pyruvate 100 mM and antibiotic-antimycotic. Macrophages were obtained by the *in vitro* differentiation of monocytes by adding 1 μM PMA to the cell culture. All cell types were grown in a humidified atmosphere at 37 °C and 5% CO_2_.

The cytotoxicity was determined by measuring cell metabolism through the Blue Cell Viability assay. Cells were seeded on MW96 microplates and incubated with the tested compounds (0.004-0.125 mg/mL) for 24 h. Control samples (not treated and chlorhexidine treated) were also analyzed. Then, the reagent was added (10%) and cells were incubated for 4 h at 37 °C. The reduction of the dye by metabolically active cells was monitored in a microplate reader (Multimode Synergy HT Microplate Reader; Biotek, USA) at 535/590 nm ex/em. Cell viability was determined by interpolation of the emission data obtained from the treated samples and the control samples (not treated cells, 100% viability).

### 4.6. Statistical analysis

Results are reported as mean ± SD. The normal distribution of the variables was analyzed by the Shapiro-Wilk test followed by the U-Mann-Whitney or Student test (StataSE 12 statistical software, StataCorp LP, USA). Statistically significant differences among groups were considered when p ≤ 0.05.

## Acknowledgements

The financial support from the Spanish Ministry of Economy and Competitiveness (grant number CTQ2014-52384-R) is gratefully acknowledged. The authors also acknowledge the financial support of the ERC Consolidator Grant program (ERC-2013-CoG-614715). CIBER-BBN is an initiative funded by the VI National R&D&i Plan 2008-2011, Iniciativa Ingenio 2010, Consolider Program, CIBER Actions and financed by the Instituto de Salud Carlos III (Spain) with assistance from the European Regional Development Fund. We acknowledge the LMA-INA and Cell Culture, Animal Care, Pathological Anatomy and Medical Imaging and Phenotyping Core Units from University of Zaragoza and IACS/IIS Aragon for their instruments and expertise. The authors declare no conflict of interest.

## References

1. Langeveld WT, Veldhuizen EJA, Burt SA. 2014. Critical Reviews in Microbiology 40:76–94

2. Hiltunen T, Virta M, Laine A-L. 2017. Philosophical Transactions of the Royal Society B-Biological Sciences 372

3. Aperce CC, Amachawadi R, Van Bibber-Krueger CL, Nagaraja TG, Scott HM, et al. 2016. Plos One 11

4. https://amr-review.org/.

5. https://www.cdc.gov/media/releases/2013/t0305lethalcre.html.

6. Laxminarayan R, Duse A, Wattal C, Zaidi AKM, Wertheim HFL, et al. 2013. The Lancet Infectious Diseases 13:1057–98

7. Chambers HF, Deleo FR. 2009. Nat. Rev. Microbiol. 7:629–41

8. Bush K, Courvalin P, Dantas G, Davies J, Eisenstein B, et al. 2011. Nat. Rev. Microbiol. 9:894–6

9. Scandorieiro S, de Camargo LC, Lancheros CAC, Yamada-Ogatta SF, Nakamura CV, et al. 2016. Frontiers in Microbiology 7

10. https://blogs.cornell.edu/cibt/files/2015/05/Antimicrobial-Functions-of-Spices.pdf.

11. Lu ZY, Chen Y, Chen W, Liu H, Song Q, et al. 2015. Journal of Antimicrobial Chemotherapy 70:653–7

12. Magi G, Marini E, Facinelli B. 2015. Frontiers in Microbiology 6

13. Walsh SE, Maillard JY, Russell AD, Catrenich CE, Charbonneau DL, Bartolo RG. 2003. Journal of Hospital Infection 55:98–107

14. Apolonio J, Faleiro ML, Miguel MG, Neto L. 2014. Fems Microbiology Letters 354:92–101

15. Becerril R, Nerin C, Gomez-Lus R. 2012. Foodborne Pathogens and Disease 9:699–705

16. Bilia AR, Guccione C, Isacchi B, Righeschi C, Firenzuoli F, Bergonzi MC. 2014. Evidence-Based Complementary and Alternative Medicine

17. Hui X, Yan G, Tian F-L, Li H, Gao W-Y. 2017. Medicinal Chemistry Research 26:442–9

18. Nostro A, Blanco AR, Cannatelli MA, Enea V, Flamini G, et al. 2004. FEMS Microbiology Letters 230:191–5

19. Cho Y, Hulang Lee. 2014. Journal of Biomedical and Translational Research 15:117–22

20. Wang L-H, Wang M-S, Zeng X-A, Zhang Z-H, Gong D-M, Huang Y-B. 2016. Journal of Agricultural and Food Chemistry 64:6355–63

21. da Luz IS, Gomes Neto NJ, Tavares AG, Nunes PC, Magnani M, de Souza EL. 2013. Archives of Microbiology 195:587–93

22. Rua J, Fernandez-Alvarez L, de Castro C, del Valle P, de Arriaga D, Garcia-Armesto MR. 2011. Foodborne Pathogens and Disease 8:149–57

23. Gavaric N, Mozina SS, Kladar N, Bozin B. 2015. Journal of Essential Oil Bearing Plants 18:1013–21

24. Ye HQ Shen SX, Xu JY, Lin SY, Yuan Y, Jones GS. 2013. Food Control 34:619–23

25. Lambert RJW, Skandamis PN, Coote PJ, Nychas GJE. 2001. Journal of Applied Microbiology 91:453–62

26. Zhou F, Ji BP, Zhang H, Jiang H, Yang ZW, et al. 2007. Journal of Food Safety 27:124–33

27. Andreu V, Mendoza G, Arruebo M, Irusta S. 2015. Materials 8:5154–93

28. Fratini F, Mancini S, Turchi B, Friscia E, Pistelli L, et al. 2017. Microbiological Research 195:11–7

29. de Sousa NTA, Gomes RC, Santos MF, Brandino HE, Martinez R, Guirro RRD. 2016. Lasers in Medical Science 31:549–56

30. Duncan B, Li X, Landis RF, Kim ST, Gupta A, et al. 2015. ACS Nano 9:7775–82

31. Ciandrini E, Campana R, Federici S, Manti A, Battistelli M, et al. 2014. Clinical Oral Investigations 18:2001–13

32. Kifer D, Muzinic V, Klaric MS. 2016. Journal of Antibiotics 69:689–96

33. Moghimi R, Aliahmadi A, Rafati H. 2017. Ultrasonics Sonochemistry 35:415–21

34. Devi KP, Sakthivel R, Nisha SA, Suganthy N, Pandian SK. 2013. Archives of Pharmacal Research 36:282–92

35. Sabulal B, Dan M, J AJ, Kurup R, Pradeep NS, et al. 2006. Phytochemistry 67:2469–73

36. Slobodnikova L, Fialova S, Hupkova H, Grancai D. 2013. Natural Product Communications 8:1747–50

37. Yuan L, Zhang YX, Jin YB, Hu YJ, Wei JY, et al. 2017. Applied Magnetic Resonance 48:201–12

38. Lv F, Liang H, Yuan QP, Li CF. 2011. Food Research International 44:3057–64

39. Kuyyakanond T, Quesnel LB. 1992. Fems Microbiology Letters 100:211–5

40. Gant VA, Warnes G, Phillips I, Savidge GF. 1993. Journal of Medical Microbiology 39:147–54

41. Di Pasqua R, Betts G, Hoskins N, Edwards M, Ercolini D, Mauriello G. 2007. Journal of Agricultural and Food Chemistry 55:4863–70

42. Xu J, Zhou F, Ji BP, Pei RS, Xu N. 2008. Letters in Applied Microbiology 47:174–9

43. Nazzaro F, Fratianni F, De Martino L, Coppola R, De Feo V. 2013. Pharmaceuticals 6:1451–74

44. La Storia A, Ercolini D, Marinello F, Di Pasqua R, Villani F, Mauriello G. 2011. Research in Microbiology 162:164–72

45. Ye H, Shen S, Xu J, Lin S, Yuan Y, Jones GS. 2013. Food Control 34:619–23

46. Krepker M, Shemesh R, Danin Poleg Y, Kashi Y, Vaxman A, Segal E. 2017. Food Control 76:117–26

47. Van den Driessche F, Brackman G, Swimberghe R, Rigole P, Coenye T. 2017. International Journal of Antimicrobial Agents 49:315–20

48. Krogsgård Nielsen C, Kjems J, Mygind T, Snabe T, Schwarz K, et al. 2017. International Journal of Food Microbiology 242:7–12

49. Xiao L, Miwa N. 2017. Human cell 30:72–87

50. Melo JO, Fachin AL, Rizo WF, Jesus HCR, Arrigoni-Blank MF, et al. 2014. Genetics and Molecular Research 13:2691–7

51. Llana-Ruiz-Cabello M, Gutierrez-Praena D, Pichardo S, Moreno FJ, Bermudez JM, et al. 2014. Food and Chemical Toxicology 64:281–90

52. Günes-Bayir A, Kiziltan H, Kocyigit A, Karatas E. A. T. 2017. Anticancer Drugs 28:522–30

53. Yu C, Liu SL, Qi MH, Zou X. 2014. Molecular Medicine Reports 9:669–76

54. https://www.drugs.com/dosage/chlorhexidine-topical.html#UsualAdultDoseforSkinDisinfectionPreoperative

55. Mendoza G, Regiel-Futyra A, Andreu V, Sebastian V, Kyziol A, et al. 2017. Acs Applied Materials & Interfaces 9:17693–701

56. Schelz Z, Molnar J, Hohmann J. 2006. Fitoterapia 77:279–85

57. Gutierrez J, Barry-Ryan C, Bourke R. 2008. International Journal of Food Microbiology 124:91–7

